# Spatial profiling of central nervous system cellular markers in murine breast cancer to brain metastases

**DOI:** 10.1101/2025.01.28.635204

**Authors:** Karen Hogg, Grant Calder, Alastair Droop, Ming Yang, Anna Simon, Mark J Hunt, Paul M Kaye, Miles Whittington, Sangeeta Chawla, William J Brackenbury

## Abstract

**Background:** Brain metastasis occurs in approximately 16% of metastatic breast cancer patients, and incidence is increasing. Patients with breast cancer brain metastases frequently develop neurological problems including cognitive impairments and seizures. We previously demonstrated accumulation of activated microglia around the tumour site in a breast cancer brain metastasis model, which co-localised with spontaneously occurring local field potential events that resembled interictal epileptic discharges. However, the mechanisms underlying these spatially restricted effects on neuronal excitability are poorly understood.

**Methods:** To better understand the underlying cellular changes and potential mechanisms, we used digital spatial profiling of brain cell subtype-specific markers to chart the spatial organisation of neurons, oligodendrocytes, astrocytes, and microglia in relation to metastatic lesion site. We developed a novel chordline-based approach to analyse these spatial changes in biomarker distribution.

**Results:** Several protein markers associated with proliferation (Ki67), astrocytes (GFAP) and microglia (CD11b, IBA1, CD45, MSR1) were upregulated in the tumour compared to healthy contralateral brain parenchyma. In contrast, some neuroglial markers (MAP2, Neurofilament light, NeuN, S100B and TMEM119) were lower in the lesion compared to normal tissue.

**Conclusions:** Overall, the protein marker changes in the lesion are indicative of neuronal cell loss and changes in the immune microenvironment of the metastatic lesion involving activated microglia, astrocytes and recruitment of peripheral immune cells. Such cellular changes may contribute to the changes in electrical activity resulting from brain metastases observed in mice and patients.

## Introduction

Brain metastasis occurs in approximately 16% of metastatic breast cancer patients, and incidence is increasing globally (Leone & Lin, 2019). Breast cancer to brain metastasis also has a poor prognosis with median survival of 10 months (Cagney *et al*., 2017). Despite recent progress, treatment options are still limited (Leone & Lin, 2019), thus this type of metastasis remains an unmet clinical need.

Patients with breast cancer brain metastases develop neurological problems including cognitive impairments (Gerstenecker *et al*., 2014) and seizures (Liede *et al*., 2023). The cellular and molecular basis of neurological symptoms is poorly understood and neurological effects are likely to be a result of non cell-autonomous mechanisms involving cancer cells, neurons, resident glial cells and infiltrating peripheral immune cells. While current treatments are targeted at reducing tumour mass and clearance of tumour cells (Bailleux *et al*., 2021), a better understanding of the cellular and molecular events that drive changes in neuronal excitability and connectivity (Sanchez-Aguilera *et al*., 2023) could help develop therapies to both clear the tumour and ameliorate neurological symptoms.

Neurological symptoms likely result from a combination of direct interactions of metastatic cells with neuronal glutamatergic signalling (Zeng *et al*., 2019) and the effects of resident glial cells and infiltrating immune cells on neuronal excitability. Glia make heterotypic contributions to metastases with individual cell types displaying pro-tumour and anti-tumor duplexity due to distinct subpopulations. For example, peritumoral astrocytes can secrete IL- 1β, which supports proliferation of metastatic cells (Mészáros *et al*., 2023); in contrast they can also deposit laminin, which promotes dormancy (Dai *et al*., 2022). Microglia, the resident brain macrophages, also have dual effects on brain metastases. Cytokines secreted from activated microglia and tumour-associated macrophages can have pro-tumour effects (Pasqualini *et al*., 2020). However, recent work suggests that pro-inflammatory microglia play an anti-tumour role in breast cancer metastasis by recruiting T cells (Evans *et al*., 2023). There is some evidence that these dual effects of microglia and astrocytes may be mediated by distinct populations with characteristic biomarkers and gene expression profile/signatures (Ishibashi & Hirata, 2024).

Our previous work in a breast cancer brain metastasis model revealed localised accumulation of activated microglia around the tumour site. We also observed spontaneously occurring local field potential events that resembled interictal epileptic discharges and with different spiking characteristics dependent on the recording position relative to the primary tumour implantation site (Simon *et al*., 2020). The sequence of events that generate these spatially distinct effects on neuronal excitability is poorly understood. To better understand the underlying cellular changes and potential mechanisms, here we used digital spatial profiling of brain cell subtype-specific markers to chart the spatial organisation of neurons, oligodendrocytes, astrocytes, and microglia. The spatial profiling panel also provided information on the distribution of activated microglia and astrocytes in relation to tumour implantation site.

## Methods

### Cell culture

MDA-MB-231 stably transduced with recombinant lentivirus for DsRed (a gift from M. Lorger, University of Leeds), PY8119 and EO771 cells (gifts from P. Ottewell, University of Sheffield) were maintained in Dulbecco’s modified eagle medium (DMEM) supplemented with 5% FBS and 4 mM L-glutamine (Simon *et al*., 2020). Cells were confirmed as mycoplasma-free using the DAPI method.

### Tumour model

Adult female wild type C57BL/6 (B6) or heterozygous *Cx3cr1*^GFP/+^ mice on a B6 background (obtained from Jackson Laboratories, Bar Harbor, USA) were used for experiments in this study. Animals were maintained on a 12 h light/dark cycle with food/water *ad libitum*. Tumour cells (10^4^ suspended in 20% v/v Matrigel) were implanted into the posterior parietal cortex following craniotomy, as described previously (Simon *et al*., 2020). Weight and body condition were monitored at regular intervals following surgery. Mice were re-grouped following recovery from surgery and were maintained for up to 1 month. Tissue was harvested for downstream analysis following perfusion fixation with 4% paraformaldehyde under terminal anaesthesia. Animal procedures were performed following approval from the University of York Animal Welfare and Ethical Review Body and under authority of a UK Home Office Project Licence. Surgical procedures were performed within the regulations of the UK Animals (Scientific Procedures) Act, 1986. Experiments have been reported in accordance with the ARRIVE guidelines.

### Tissue preparation and immunohistochemistry

Brains were placed in fixative at 4°C for 48 h, cryopreserved in 30% sucrose solution for 24 h, and then frozen in OCT blocks and stored at -70°C. Coronal sections (5 µm thick) were cut onto Superfrost+ slides using a Leica cryostat and either processed for digital spatial profiling or conventional immunostaining. For GFAP staining, slices on slides were permeabilised in PBS containing 0.5% Triton-X for 8 minutes followed by a further 2 minute incubation in PBS containing 0.2% Triton-X and 10% methanol. Slices were blocked with PBS containing 3% normal goat serum for 3 h and then incubated overnight at 4°C with a rabbit GFAP antibody (Dako Z0334) diluted 1:300 in PBS containing 0.2% Triton X100 and 1% normal goat serum. Alexa Fluor 568-conjugated anti-rabbit antibody (made in goat; ThermoFisher Scientific A-11011) was used at 1:500 dilution in PBS containing 1% normal goat serum and sections were counterstained with Hoechst 33258.

Overview images of the brain slices were collected using a Zeiss LSM980 microscope set to standard epi-fluorescent mode using an AxioCam 305 mono camera. Images were captured with a 20x/0.8 air objective lens and the field of view expanded to cover the whole tissue using image tiling. The overlapping images (10%) were stitched into a single image using Zen Blue “stitch”. The fluorescent dyes excitation and emission were captured at the following wavelengths: Hoechst 33258 was excited at 370–410 nm and the emission collected at 430–470 nm; Alexa Fluor 488 was excited at 450–490 nm and emission collected at 500–550 nm; Alexa Fluor 568 was excited at 538–563 nm and emission collected at 570–640 nm.

Tissue subregions were selected for higher detail examination using confocal modality. Images were captured with a 40x/1.3 oil immersion objective lens and the area expanded using 2×2 tile. To obtain all detail throughout the tissue section, Z stacks with a step interval 0.27 µm were collected. Z-stack images were maximum intensity projected using Zen Blue “Extended focus” tool. Hoechst 33258 was excited using 405 nm laser line and emission collected at 403–489 nm; Alexa Fluor 488 excited using 488 nm laser line and emission collected 493–553 nm and Alexa Fluor 568 excited 561 nm laser line and emission collected 575–757 nm.

### Digital spatial profiling

Spatial profiling of cryosections were performed using a GeoMX Digital Spatial Profiler (NanoString), following the manufacturer’s established guidelines. For detailed sample preparation protocol, see “GeoMx DSP Manual Slide Preparation User Manual” section “Protein Slide Preparation Protocol (FFPE)” with the modification in Appendix II - “Fresh frozen sample preparation for protein assays” (NanoString, n.d.).

Cryosections were fixed in 10% NBF overnight (12–16 hours) at room temperature. To ensure compatibility with the GeoMx antibody panels, antigen retrieval was performed using Citrate buffer pH 6.0 in a pressure cooker set to “high pressure and temperature” for 15 minutes. The samples were blocked using GeoMx Buffer W for 1 hour at room temperature. Cryosections were then stained overnight at 4°C with both a marker to define the relevant tissue compartments (anti-GFP to identify CX3CR1-GFP+ microglia) plus the GeoMx Neuronal Cell profiling & Glial cell subtyping panels. Excess antibody probes were washed off and samples were post-fixed using 4% PFA for 30 minutes. An additional morphology marker, SYTO 83 was used to stain nuclei to aid tissue detection. Samples were mounted in the GeoMx DSP system where overview images of the tissue were captured using the morphology markers. These overview images were then used to determine the areas of the tissue to be analysed by the targeted release of UV photocleavable oligonucleotide barcodes conjugated to the GeoMx antibodies targeting 29 neuroglial markers, three housekeepers (histone H3, ribosomal protein S6 and GAPDH) and three negative controls (rabbit IgG, rat IgG2a and rat IgG2b) (Supplementary Table 1). The identifying barcodes that were released were identified and counted using the NanoString nCounter system. For each animal, four regions of interest (ROIs) were drawn (repeated across at least three tissue sections, to give a total of at least 12 ROIs per mouse). The ROIs were defined as follows:

1. “Lesion” – a freeform ROI across the tumour area, as defined by GFP and/or SYTO 83 staining.
2. “Distal” parenchyma – a circle (250 µm in diameter), 100 µm from edge of lesion ROI.
3. “Far” parenchyma – a circle (250 µm in diameter), 100 µm from the nearside edge of the distal ROI.
4. “Contralateral” – a freeform ROI of the same shape and size as the lesion, placed on the contralateral side of the brain.

Protein expression counts were normalised to internal spike-in hybridisation controls and then normalised to the geometric mean of the three housekeepers within each ROI.

### Data processing and statistical analysis

The results from the nCounter readout were loaded back into the GeoMx DSP data center software for further analysis. Quality Control of the data was assessed by analysing the internal/spiked in hybridisation controls to ensure the integrity of the nCounter readout. Further, positive control (housekeeping proteins) and negative control (isotype) antibody probes were used to assess antibody labelling of the sample and for normalisation steps. After initial QC assessment, data were exported for further analysis in R using the standR package (Liu et al., 2023). Differential analysis was performed using LIMMA (Ritchie *et al*., 2015). In order to succinctly show the behaviour of individual proteins between the different distances from the original lesion, we generated per-protein normalised intensity plots (termed chordline plots) as follows: the total range of expression values at each location (Lesion, Distal, Far and Contralateral) across all selected proteins was calculated, and the resulting range split into 5 equal sections. The value for a single protein was then assigned into one of these categories to generate a 4-value “chordline”. Scripts are available on Github at https://github.com/uoy-research/2024_spatial_profiling_brain_mets.

## Results

### Study cohort

The study cohort is described in Table 1. Matched ROIs (comprising Lesion, Distal, Far and Contralateral) were evaluated from a total of five female mice. All mice were on the B6 background, and four were *Cx3cr1*^GFP/+^. Three mice were implanted with MDA-MB-231 cells, one mouse with EO771 cells and one mouse with PY8119 cells. An example coronal brain section from a *Cx3cr1*^GFP/+^ mouse implanted with MDA-MB-231 cells highlighting the four ROIs is in Figure 1A. There was considerable variation in expression of the 29 neuroglial markers between the lesion and the distal, far and contralateral ROIs across all five mice (Figure 1B). Principal component analysis revealed an absence of clustering by implanted cell line or mouse (Supplementary Figure 1A, B). Similarly, unsupervised hierarchical clustering of protein expression by all factors included in the study revealed no obvious patterns relating to cell line, mouse, slide, or marker annotation class (Supplementary Figure 2).

**Table 1.** Sample cohort characteristics.

| Characteristic | n | % |
| --- | --- | --- |
| Mice | 5 | 100 |
| Tissue sections | 9 | 100 |
| Slides | 5 | 100 |
| Regions of interest |  |  |
| Lesion | 26 | 27 |
| Distal | 22 | 23 |
| Far | 22 | 23 |
| Contralateral | 26 | 27 |
| Genotypes |  |  |
| Wildtype | 1 | 25 |
| *Cx3cr1*<sup>GFP/+</sup> | 4 | 75 |
| Sex – female | 5 | 100 |
| Cell line implants (mice) |  |  |
| MDA-MB-231 | 3 | 60 |
| EO771 | 1 | 20 |
| PY8119 | 1 | 20 |

**Figure 1.**
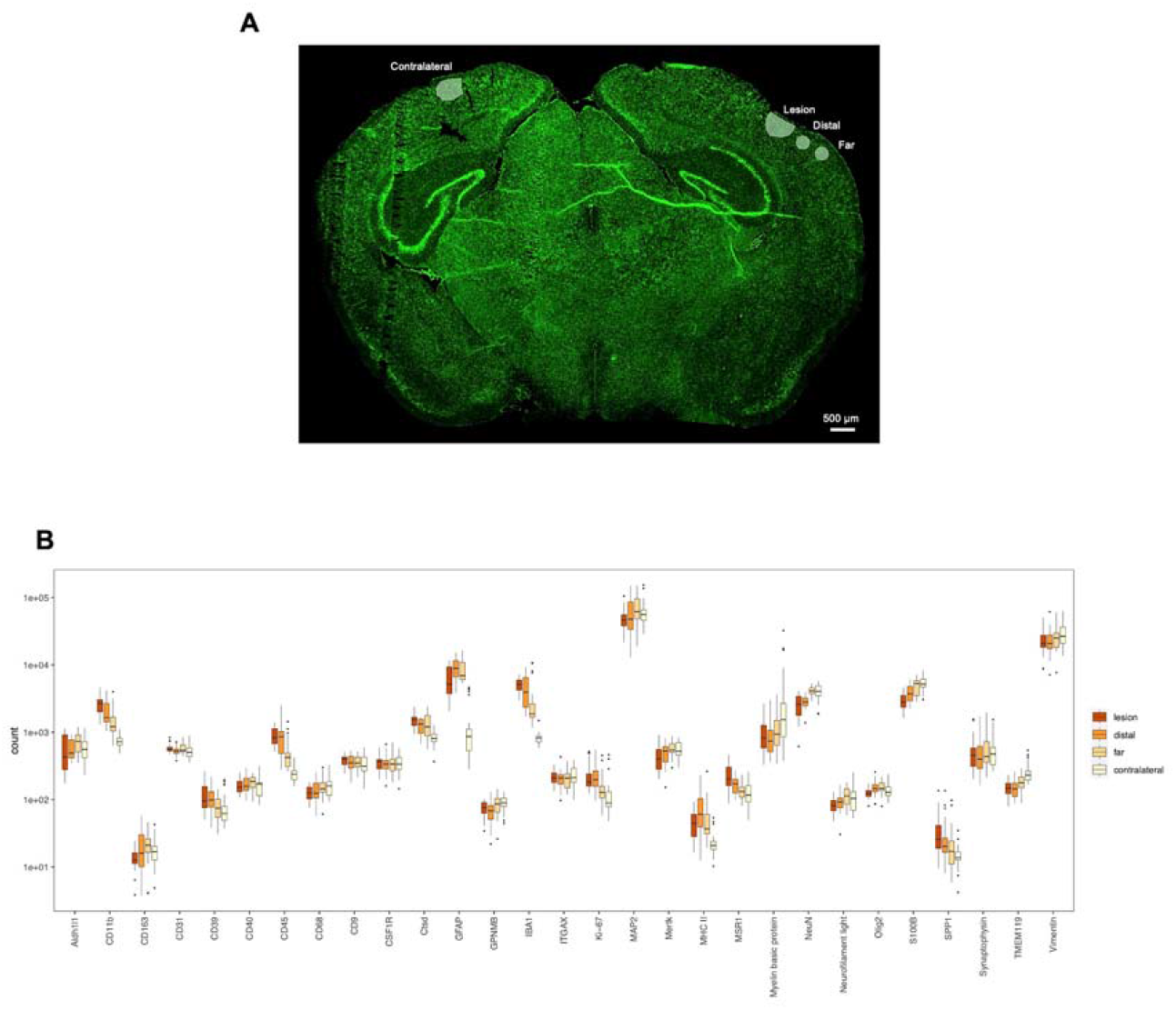
Neuronal cell profiling marker protein expression heterogeneity across metastatic lesions and adjacent brain parenchyma. (A) Example image of coronal brain section highlighting lesion, distal, far and contralateral region of interest (ROI) locations. Anti-GFP staining in green indicates CX3CR1-GFP+ microglia. (B) Box plot comparing marker expression (normalised to housekeeper proteins) for each ROI location. Boxes extend from the 25th to the 75th percentile, with the median displayed as a bold line. Whiskers extend to the furthest values within 1.5 × IQR of the box. Dots show any points outside this range. n = 22–26 ROIs per location, pooled from a total of 5 mice.

### Region-specific heterogeneity

We next compared expression of neuroglial protein markers between specific ROI locations. We initially compared the lesion with the normal brain parenchyma on the contralateral side (Figure 2A). This comparison revealed a significant upregulation of several markers in the lesion which are associated with accumulation of astrocytes (GFAP) and microglia (CD11b, IBA1, CD45, MSR1), as well as increased cell proliferation (Ki67) (Figure 2A; Supplementary Table 1). Next, we compared the lesion with the ipsilateral far ROI containing nearby histologically normal brain parenchyma (Figure 2B). This second comparison revealed fewer markers significantly upregulated in the lesion: MSR1 and CD45, both associated with accumulation of microglia. Notably, no neuroglial protein markers were significantly downregulated in the lesion ROI compared to ipsilateral far or contralateral ROIs (Figure 2A, B). In support of the general upregulation of astrogliosis and proliferation markers in the lesion when compared with the contralateral brain parenchyma, principal component analysis revealed marked clustering of lesion versus contralateral ROIs (Figure 2C). However, there was more overlap between lesion and ipsilateral distal and far ROIs, consistent with the tumour potentially affecting microglial activation several hundreds of microns away from the lesion margin.

**Figure 2.**
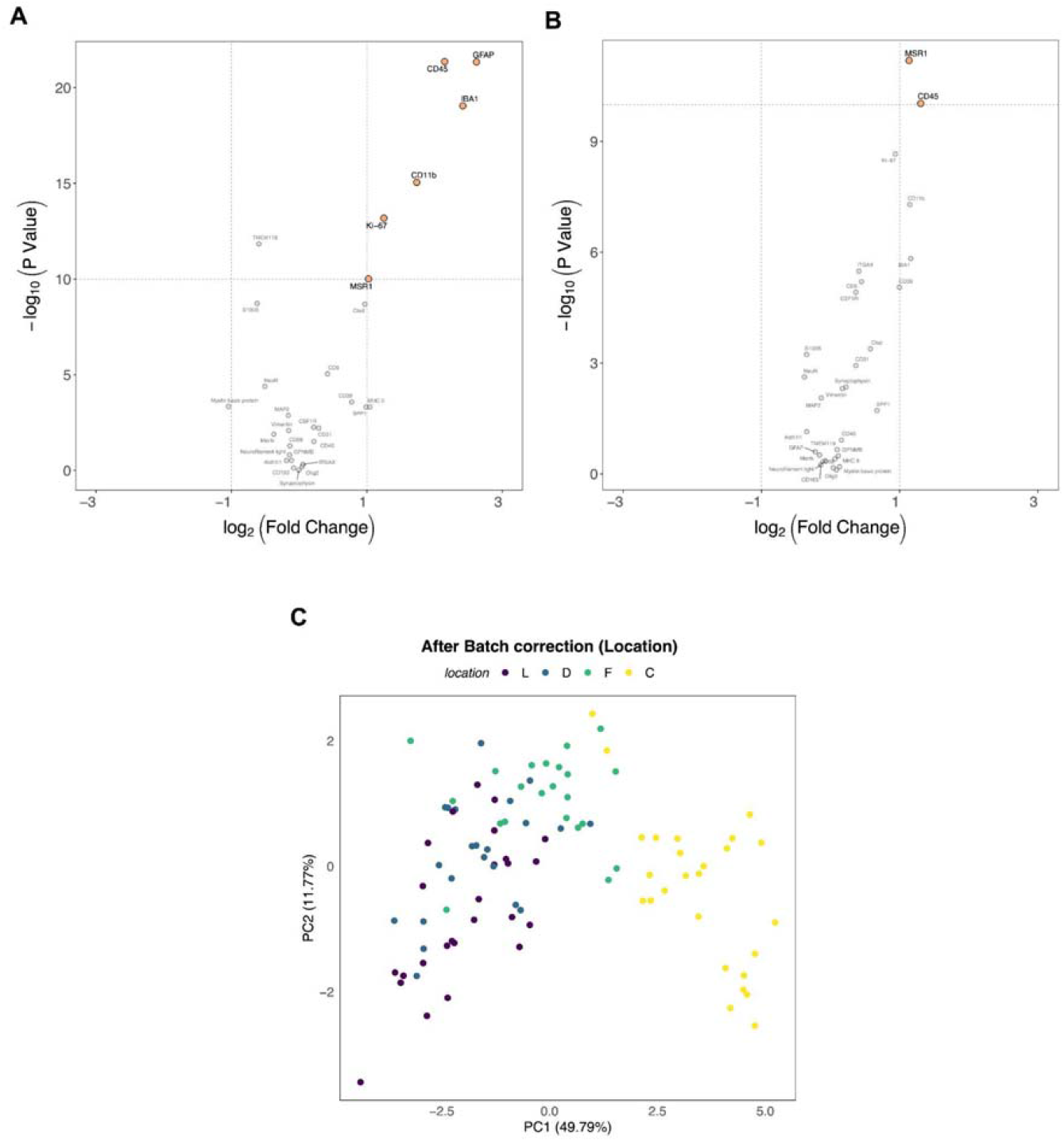
Differential neuronal cell profiling marker protein expression by tissue compartment. (A) Volcano plot comparing marker expression between lesion (right) versus contralateral (left) region of interest (ROI) locations. (B) Volcano plot comparing marker expression between lesion (right) versus far (left) ROI locations. Statistical analysis was performed using LIMMA and included false discovery rate (FDR) correction for multiple comparisons. (C) Principal component analysis (PCA) biplot of marker protein expression within each ROI, with locations highlighted by colour. n = 96 ROIs from a total of 5 mice.

### Marker differences across regions

In order to visualise key patterns in expression levels of protein markers across the different ROI regions, and relate these to distance from the tumour, we generated a per-protein normalised intensity (chordline) plot (Figure 3). The chordline plot showed general upregulation of several proliferation, microglial and astrocyte markers towards the lesion, confirming the results from Figure 2. However, some other astrocyte and microglial markers (S100B, TMEM119) were lower in the lesion than in the normal tissue. In addition, neuronal markers (MAP2, Neurofilament light, NeuN) were lower in the lesion, gradually increasing through distal and far regions to the contralateral side, suggesting a general loss of neuronal density/viability within and around the lesion. Moreover, GFAP expression, while higher in the lesion compared to the contralateral side, was upregulated further in the distal and far ROIs, suggesting possible accumulation of astrocytes surrounding and infiltrating the lesion. Broadly similar patterns were evident in chordline plots separated by implanted cell type (Supplementary Figures 3 & 4).

**Figure 3.**
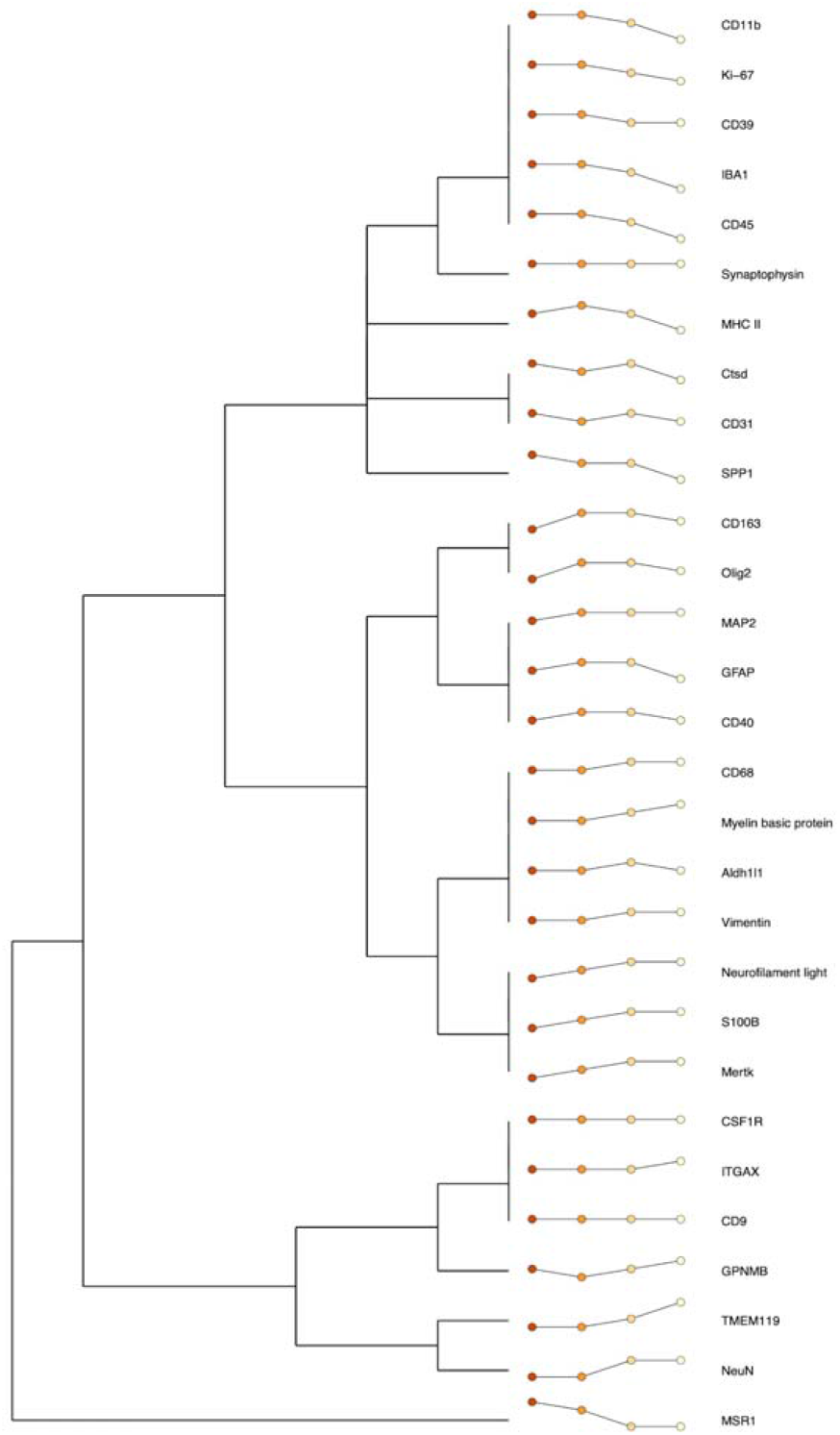
Neuronal cell profiling marker protein expression across region of interest (ROI) location represented as chordlines. Per-protein normalised intensity plots of expression at each location (left to right: Lesion (red), Distal (orange), Far (yellow) and Contralateral (cream)) with range split into 5 equal sections to generate a 4-value chordline.

To further validate the apparent astrocyte accumulation near to and within the lesion, we next studied GFAP distribution using immunohistochemistry. Confocal microscopy revealed notable accumulation of activated microglia in the lesion, in agreement with previous observations (Simon *et al*., 2020) (Figure 4A–C). Interestingly, accumulation of GFAP+ astrocytes was even more marked, and extended beyond the immediate lesion into nearby ipsilateral brain parenchyma (Figure 4A–C). In contrast, GFAP+ astrocyte distribution was unaltered in the ipsilateral vs contralateral hippocampus (Supplementary Figure 5). Together, these spatial analyses of neuroglial protein marker distribution support a considerable accumulation of both activated microglia and astrocytes both within, and surrounding metastatic breast cancer brain lesions.

**Figure 4.**
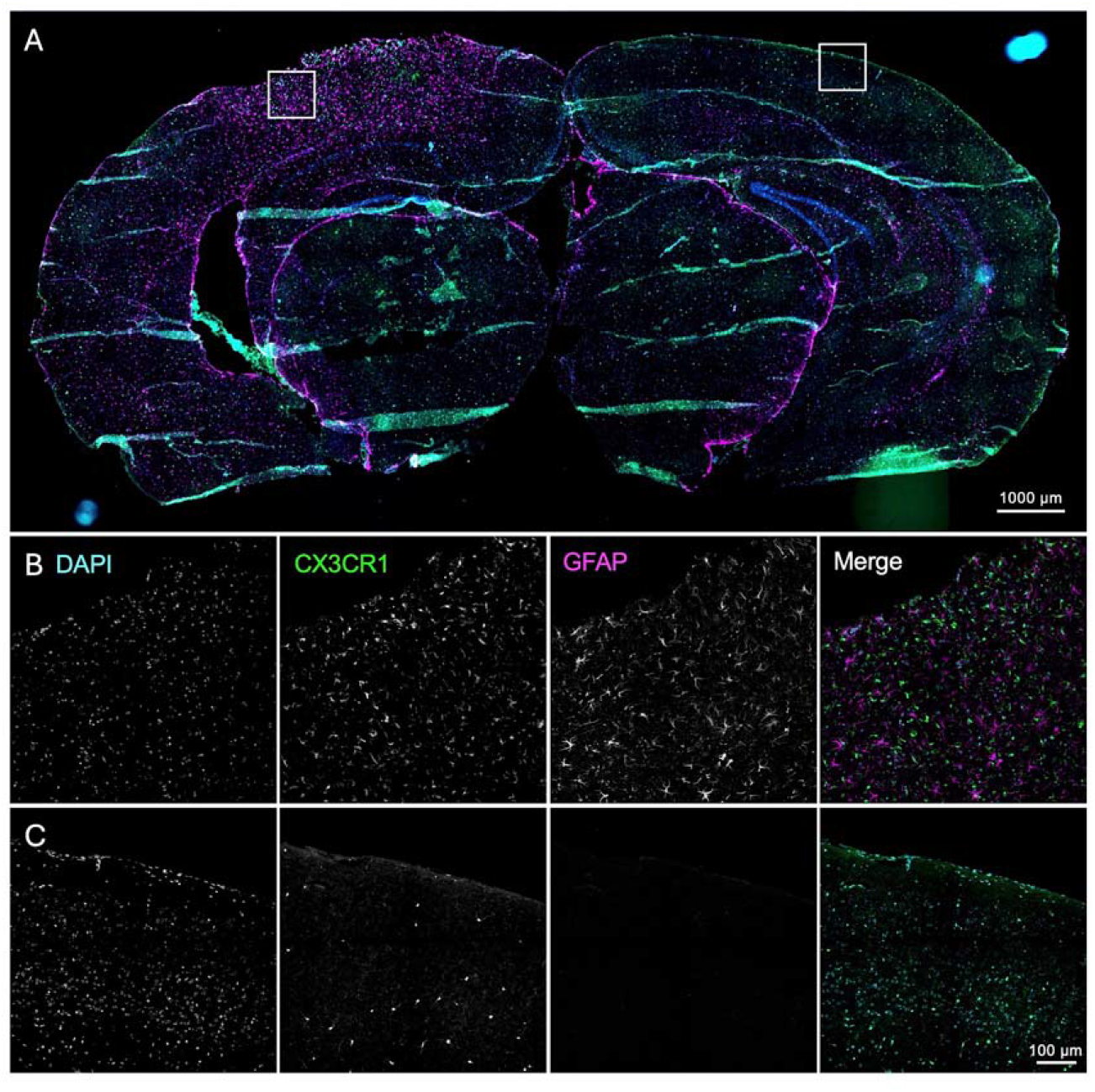
Accumulation of astrocytes near the lesion. Confocal images of CX3CR1-GFP+ microglia (green), GFAP+ astrocytes (magenta) and nuclei (blue) in a coronal section obtained from a Cx3cr1GFP/+ mouse euthanised 6 days following implantation of MDA-MB-231 cells. (A) Overview image of whole section. (B) Expanded ROI within lesion. (C) Expanded ROI on the contralateral side.

## Discussion

In this study, we used digital spatial profiling to understand the distribution of neurons, oligodendrocytes, astrocytes and microglia within and adjacent to lesions in a mouse model of breast cancer brain metastasis. Our findings show that several protein markers associated with proliferation (Ki67), astrocytes (GFAP) and microglia (CD11b, IBA1, CD45, MSR1) were upregulated in the tumour compared to healthy contralateral brain parenchyma. Fewer proteins were upregulated in the lesion when compared to ipsilateral normal parenchyma. These observations were supported by principal component analysis revealing clustering of lesion versus contralateral tissue, with further overlap between lesion and ipsilateral tissue. Interestingly, some neuroglial markers (MAP2, Neurofilament light, NeuN, S100B and TMEM119) were lower in the lesion compared to normal tissue. In agreement with our data, TMEM119 expression has been shown to decrease in Alzheimer’s disease-associated microglia, along with other microglial ‘homeostatic genes’ (Keren-Shaul *et al*., 2017; Kenkhuis *et al*., 2022), supporting the notion that the metastatic lesion leads to remodelling of microglial function. Immunohistochemical analysis further supported the observation that CX3CR1-GFP+ microglia and GFAP+ astrocytes accumulated within, and adjacent to, the lesion. Overall, the protein marker changes in the lesion are indicative of neuronal cell loss and changes in the immune microenvironment of the metastatic lesion involving activated microglia, astrocytes, and possibly other peripheral immune cell types.

Tumorigenic interactions involving multiple cell types in the brain metastatic niche could explain the pattern of electrical changes we have previously reported (Simon *et al*., 2020). To better understand the relationship between cortical electrical activity and brain metastasis, we utilised a novel chordline-based approach to track protein marker changes within the lesion, where electrical activity changes are prominent, and across distal and far areas displaying normal electrical activity. This approach allowed us to identify how protein markers, and thus cell types, are spatially clustered. We observed several distinct patterns of particular sets of markers. Of note, the CD11b, Ki-67, CD39, IBA1 and CD45, indicative of inflammation via microglial activation and recruitment of peripheral macrophages, all showed highest expression in the lesion with a progressive decrease moving away from the primary lesion site. Markers for activated astrocytes showed higher expression further away from the lesion, with GFAP, Aldh1 and vimentin all being higher in peritumoral regions. This pattern of indicative astrogliosis is similar to that seen in ischemic stroke models where GFAP immunoreactivity is highest in the peri-infarct regions (Noll *et al*., 2022). Moreover, in the ischemic stroke model, the core border showed extensive immune activation markers, similar to what we observed in the lesion site region.

The spontaneous multi-unit activity within the lesion and field spike activity at the border of the lesion that we observed previously (Simon *et al*., 2020) may be driven by inflammatory mediators underpinned by microglial accumulation and/or activation. The unique spatially resolved molecular signature identified in this study may also highlight altered astrocyte function or astrogliosis, in addition to microglia/macrophage activation. It is noteworthy that the astrocyte-secreted protein S100B is lower in the lesion and peritumoral areas compared to the contralateral side, given that S100B knockout mice display increased epileptiform activity (Dyck *et al*., 2002). Interestingly, the spatial pattern of S100B expression is inverse to that of GFAP, and has been reported before in a viral infection model (Ohtaki *et al*., 2007). This inverse relationship suggests heterogeneous astrocyte function as a result of the metastatic lesion. Indeed, both markers have independent characteristics and may yield complementary information relating to astrocyte function (Janigro *et al*., 2022).

In conclusion, we developed a novel approach to evaluate the spatial pattern of neuroglial biomarker expression across brain tissue using chordlines. This approach reveals unique cellular differences between the lesion site and peritumoral areas, and highlights opportunities to develop more targeted mechanistic investigations and therapeutic approaches with the goal of ameliorating electrophysiological sequelae of brain metastasis.

## Declarations

## Ethics approval and consent to participate

Animal procedures were performed following approval from the University of York Animal Welfare and Ethical Review Body and under authority of a UK Home Office Project Licence. Surgical procedures were performed within the regulations of the UK Animals (Scientific Procedures) Act, 1986.

## Consent for publication

Not applicable.

## Availability of data and materials

Scripts are available on Github at https://github.com/uoy-research/2024_spatial_profiling_brain_mets. The datasets used and/or analysed during the current study are available from the corresponding author on reasonable request.

## Authors’ contributions

KH, GC, AD, MY, PK, MW, SC and WB contributed to the conception and design of the work. KH, GC, AD, MY, AS, MH, PK, MW, SC and WB contributed to acquisition, analysis, and interpretation of data for the work. KH, GC, AD, PK, SC and WB contributed to drafting the work and revising it critically for important intellectual content. All authors approved the final version of the manuscript.

## Acknowledgements

The authors gratefully acknowledge assistance from Genomics Lab in the Bioscience Technology Facility at the University of York. This paper is dedicated to Professor Miles Whittington who sadly passed away in 2021.

