## Supplementary Data for "Spatial profiling of central nervous system cellular markers in murine breast cancer to brain metastases"

**Supplementary Table 1. Protein marker metadata.**

| Class | Name | Annotations |
|---|---|---|
| Housekeeping control | Histone H3 | NA |
| Housekeeping control | GAPDH | NA |
| Housekeeping control | Ribosomal protein S6 | NA |
| Negative control | Rb IgG | NA |
| Negative control | Rt IgG2a | NA |
| Negative control | Rt IgG2b | NA |
| Endogenous target | Aldh1l1 | Astrocyte |
| Endogenous target | CD11b | Microglia |
| Endogenous target | CD163 | Microglia |
| Endogenous target | CD31 | Endothelia |
| Endogenous target | CD39 | Inflammation; Microglia |
| Endogenous target | CD40 | Inflammation; Microglia |
| Endogenous target | CD45 | Inflammation; Microglia |
| Endogenous target | CD68 | Microglia |
| Endogenous target | CD9 | Disease-Associated Microglia |
| Endogenous target | CSF1R | Microglia |
| Endogenous target | Ctsd | Microglia |
| Endogenous target | GFAP | Inflammation; Astrocyte |
| Endogenous target | GPNMB | Disease-Associated Microglia |
| Endogenous target | IBA1 | Microglia |
| Endogenous target | ITGAX | Microglia |
| Endogenous target | Ki-67 | Proliferation |
| Endogenous target | MAP2 | Neuron; Cytoskeleton |
| Endogenous target | Mertk | Disease-Associated Microglia |
| Endogenous target | MHC II | Microglia |
| Endogenous target | MSR1 | Microglia |
| Endogenous target | Myelin basic protein | Oligodendrocytes |

| | | |
|---|---|---|
| Endogenous target | NeuN | Neuron |
| Endogenous target | Neurofilament light | Neuron; Cytoskeleton |
| Endogenous target | Olig2 | Oligodendrocytes |
| Endogenous target | SPP1 | Disease-Associated Microglia |
| Endogenous target | Synaptophysin | Synaptic Vesicle |
| Endogenous target | S100B | Astrocyte |
| Endogenous target | TMEM119 | Microglia |
| Endogenous target | Vimentin | Astrocyte |

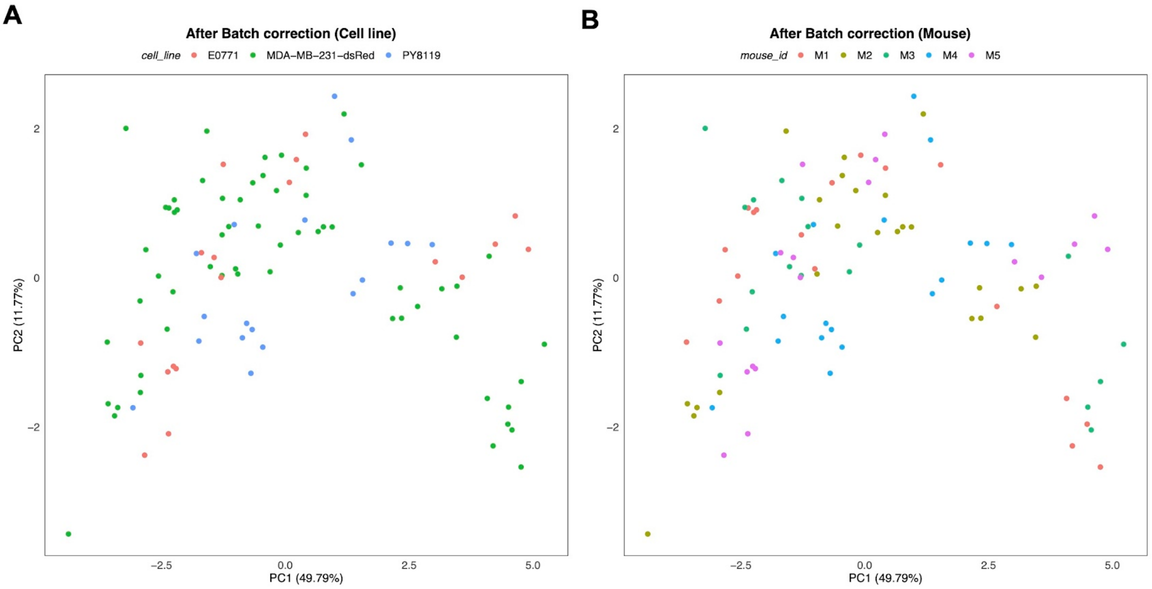


**Supplementary Figure 1. Principal component analysis (PCA) biplots.**

(A) PCA biplot of marker protein expression within each ROI, with implanted cell line highlighted by colour. (B) PCA biplot of marker protein expression within each ROI, with recipient mice highlighted by colour. n = 96 ROIs from a total of 5 mice.

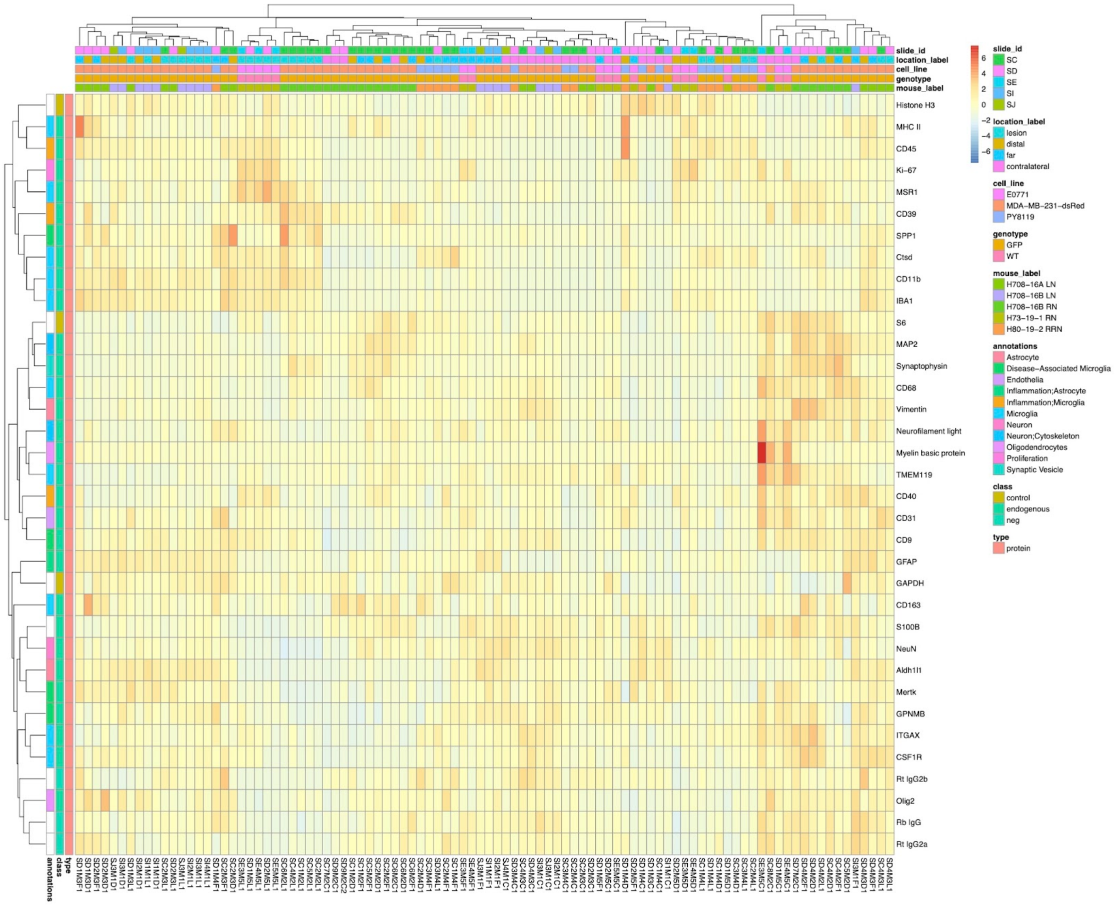


**Supplementary Figure 2. Heatmap of neuronal cell profiling marker protein expression level clustered by region of interest (ROI) location, cell line, recipient mouse, slide ID, and marker annotation class.**

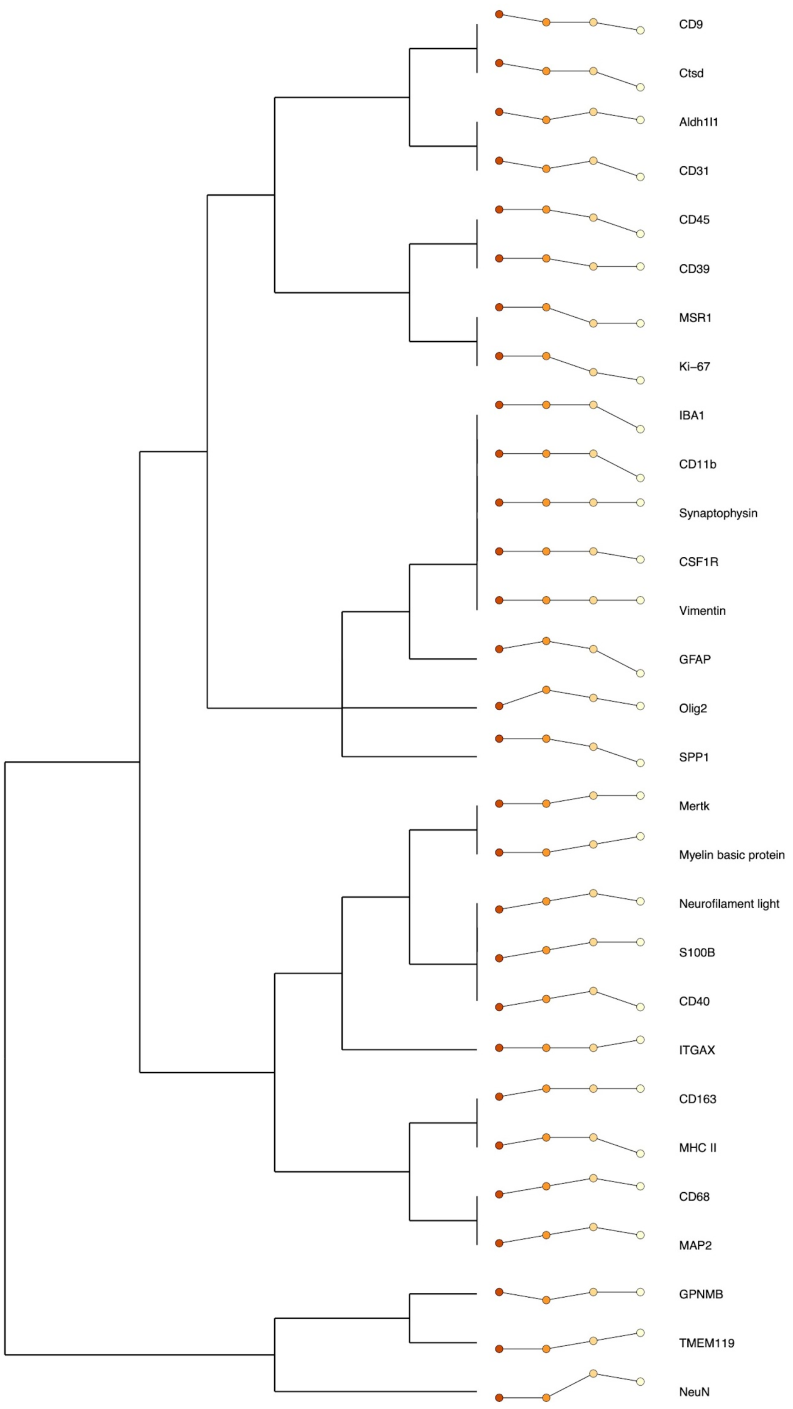


**Supplementary Figure 3. Neuronal cell profiling marker protein expression across region of interest (ROI) location represented as chordlines in MDA-MB-231 xenografts.** Per-protein normalised intensity plots of expression at each location (left to right: Lesion, Distal, Far and Contralateral) with range split into 5 equal sections to generate a 4-value chordline.

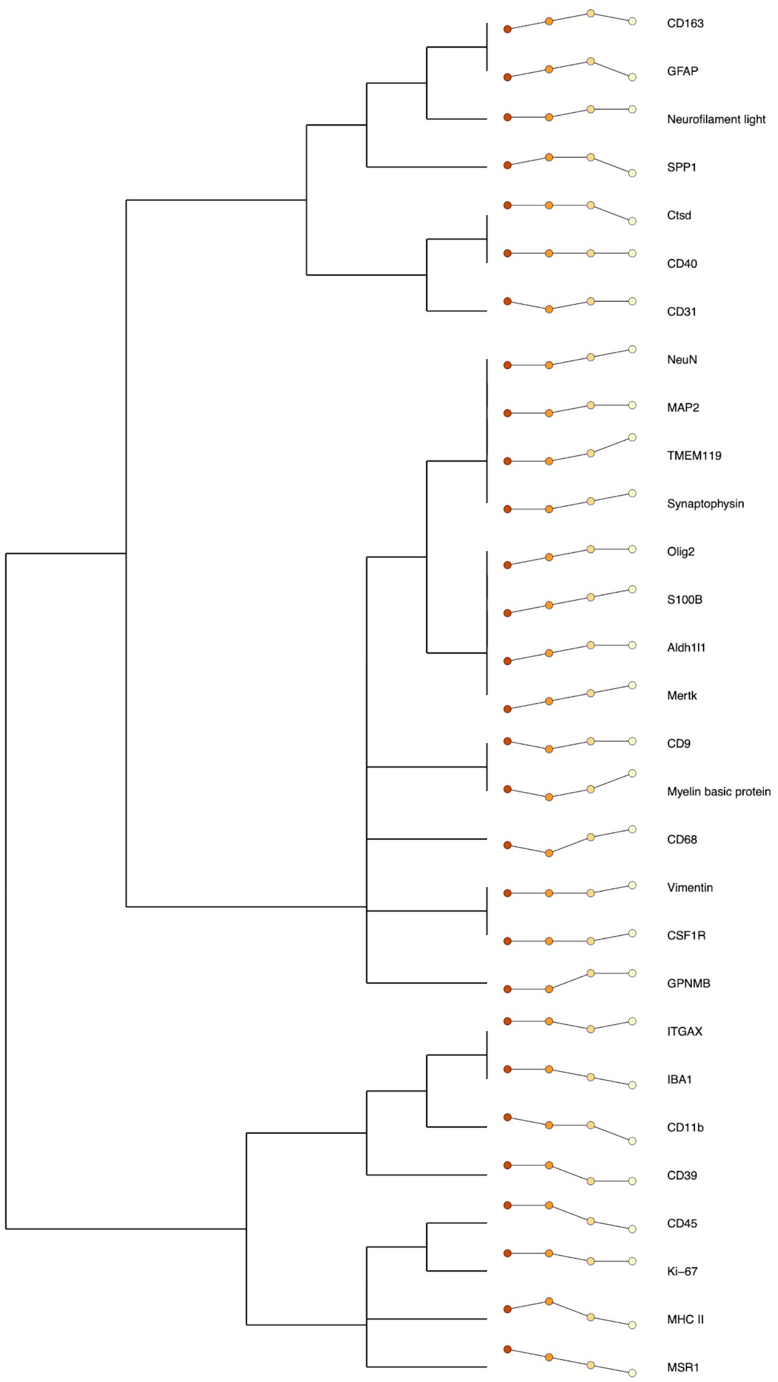


**Supplementary Figure 4. Neuronal cell profiling marker protein expression across region of interest (ROI) location represented as chordlines in EO771/PY8119 allografts.**

Per-protein normalised intensity plots of expression at each location (left to right: Lesion, Distal, Far and Contralateral) with range split into 5 equal sections to generate a 4-value chordline.

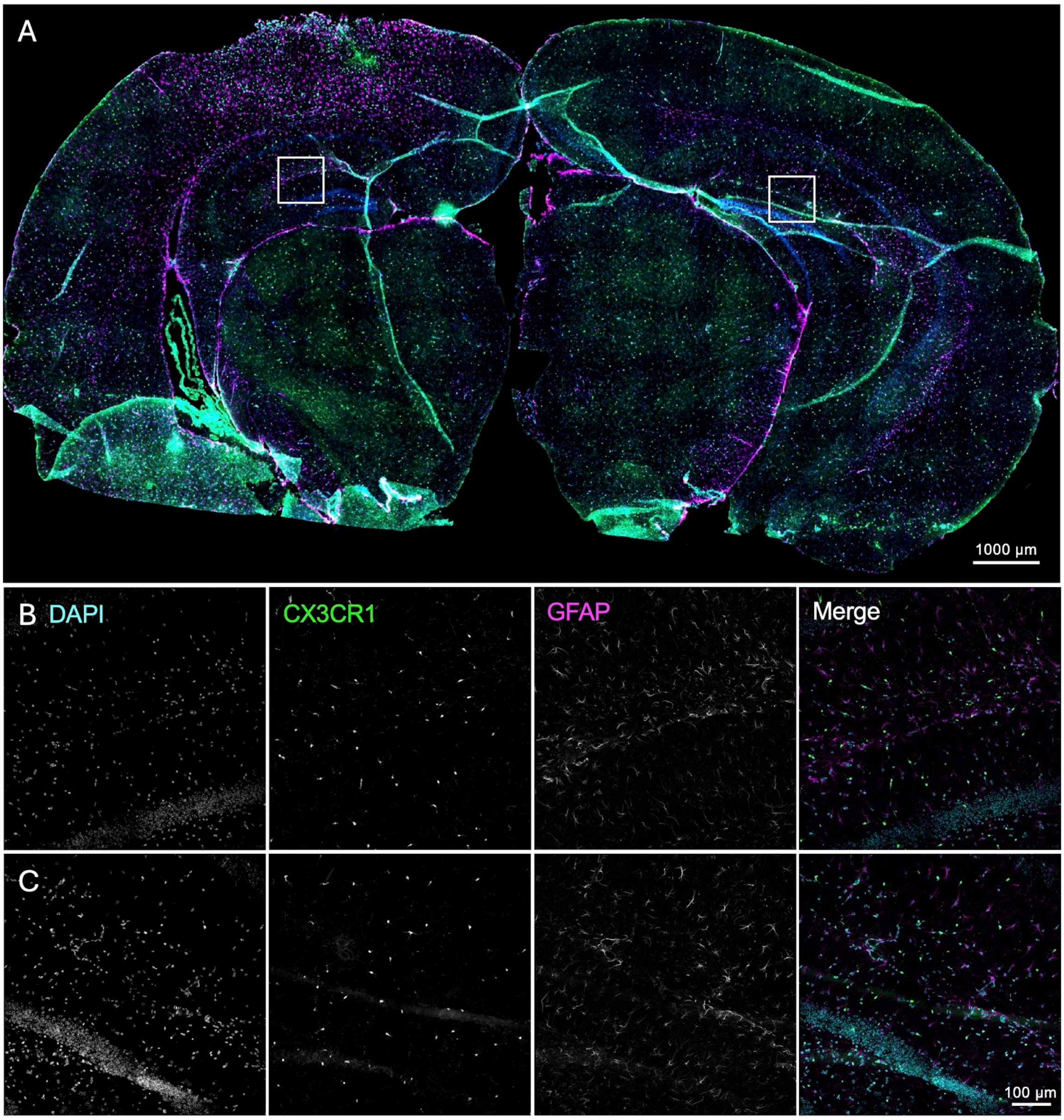


**Supplementary Figure 5. Astrocyte distribution is unaltered in the hippocampus.** Confocal images of CX3CR1-GFP+ microglia (green), GFAP+ astrocytes (magenta) and nuclei (blue) in a coronal section obtained from a Cx3cr1GFP/+ mouse euthanised 6 days following implantation of MDA-MB-231 cells. (A) Overview image of whole section. (B) Expanded ROI in the hippocampus on the ipsilateral side to the lesion. (C) Expanded ROI in the hippocampus on the contralateral side to the lesion.
